# Parametric and Non-parametric Gradient Matching for Network Inference

**DOI:** 10.1101/254003

**Authors:** Leander Dony, Fei He, Michael PH Stumpf

## Abstract

Reverse engineering of gene regulatory networks from time series gene-expression data is a challenging problem, not only because of the vast sets of candidate interactions but also due to the stochastic nature of gene expression. To avoid the computational cost of large-scale simulations, a two-step Gaussian process interpolation based gradient matching approach has been proposed to solve differential equations approximately. Based on this gradient matching approach, we evaluate the fits of parametric as well as non-parametric candidate models to the data under various settings for different inference objectives. We also use model averaging, based on the Bayesian Information Criterion (BIC), in order to combine the different inferences. We found that parametric methods can provide comparable, and often improved inference compared to non-parametric methods; the latter, however, require no kinetic information and are computationally more efficient.

The code used in this work is available at https://github.com/ld2113/Final-Project.

## 1 Introduction

Gene expression is known to be subject to sophisticated and fine-grained regulation. Besides underlying the developmental processes and morphogenesis of every multicellular organism, gene regulation represents an integral component of cellular operation by allowing for adaptation to new environments through protein expression on demand [1, 2, 3, 4].

While the basic principles of gene regulation have been discovered as early as 1961 [5], understanding the structure and dynamics of complex gene regulatory networks (GRN) remains an open challenge. In this context experimental analysis has been complemented by a rich and growing literature on network reconstruction or inference, where statistical tools are used to “learn” or reconstruct interactions between sets of genes from data [1, 6, 7, 8].

Given the vast range of network inference approaches studied within and outside the life sciences, we necessarily have to limit our analysis, and we focus on a small but diverse set of inference approaches. Further details on other methods can be found in references [9, 10].

Gene regulatory interactions within a group of genes can be visualised in various ways. Usually, genes and their interactions are represented as nodes and edges of a graph respectively. Depending on the aim of the study and the employed method, the graph can be undirected (Figure 1A); directed (Figure 1B); or contain further information about interaction types (Figure 1C).

When inferring candidate regulatory networks from time-course data (e.g. time-resolved mRNA concentration measurements), different mathematical representations of the individual interactions can be used. The application of ordinary differential equation (ODE) models in this context has the advantage that each individual term in the final ODE model can provide direct mechanistic insight (such as presence of activation or repression) [11, 12]. Following [13] the ODEs of a GRN can e.g. be expressed as,

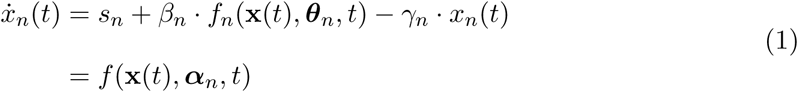

Here, *x*_*n*_(*t*) denotes the concentration of *n*^*th*^ mRNA at time *t*, *s*_*n*_ is the basal transcription rate, γ_n_ is the mRNA decay rate, x is a vector of concentrations of all the parent mRNAs that regulate the *n*^*th*^ mRNA, the regulation function *f*_*n*_ describes the regulatory interactions among genes such as activation or repression that are normally quantified by Hill kinetics, with *β*_*n*_ the strength or sensitivity of gene regulation, and the parameter vector *θ*_n_ contains regulatory kinetic parameters. The right-hand-side of the *n*^*th*^ ODE can be summarized in a single nonlinear function *f* with *α_n_* including all the kinetic parameters. Some approaches such as non-parametric Bayesian inference methods provide less mechanistic information but they may nevertheless provide realistic representations of complex regulatory interactions between genes, which a simple ODE system might not be able to capture [14], especially when accurate kinetic information is unavailable.

**Figure 1:**
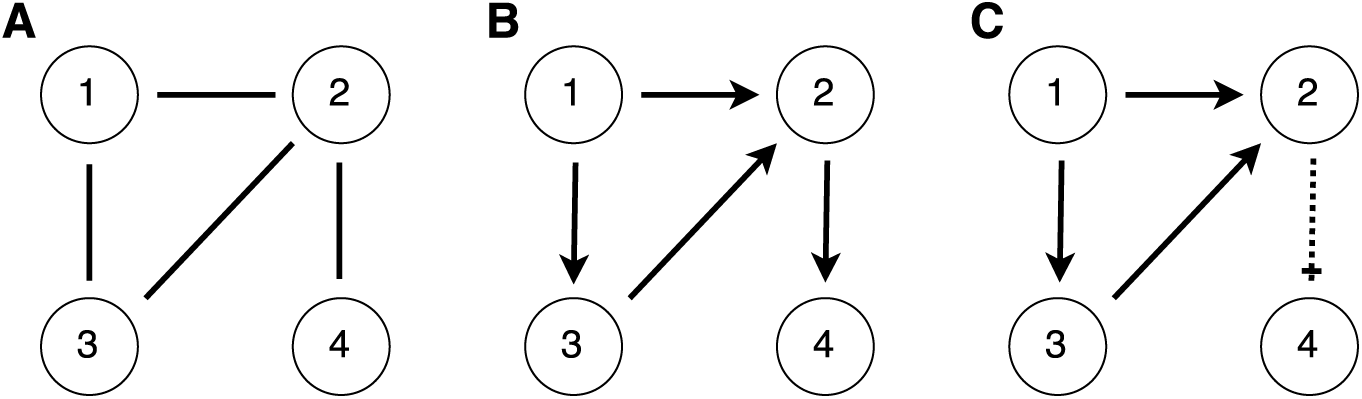
Gene regulatory network (GRN) schematics with four genes and four interactions. Three representations of the same GRN are shown. A Undirected graph showing interactions between genes: 1, 2; 1, 3; 2, 3; 2, 4. B Directed graph showing interactions between genes (parent node stated first): 1, 2; 1, 3; 3, 2; 2, 4. C Directed graph showing interactions between genes: 1 activates 2; 1 activates 3; 3 activates 2; 2 represses 4.

Parameter and structure inference of a mathematical model expressed as coupled ODEs (1) is a challenging problem, as repeatedly solving the ODEs by numerical integration is required which is computationally costly. Such costs quickly increase as the number of genes in the network increases. A two-step gradient matching approach has been proposed in the machine learning literature [15, 16, 17] to reduce the computational cost: in the first step, the time series data are interpolated, and in the second step, the parameters of ODEs are optimized by minimizing the difference between interpolated derivatives and the right-hand-side of ODEs. Thus the ODEs do not need to be solved explicitly. Previous work in the field of automatic network reconstruction has proposed a gradient matching approach to triaging different network topologies [11, 18]. Additionally, gradient matching enables treating each species in the network independently.

Thus gradient matching for automatic ODE network reconstruction combined with Gaussian process (GP) regression could be a promising avenue for inferring GRNs. But still, some problems remain: model identifiability, as too many models provide a good fit to the data; reliably fitting HGPs to noisy data; and potentially limiting model assumptions, e.g. by considering only a limited range of interaction types.

In this work, we investigate and attempt to address those issues and furthermore evaluate inference performance of gradient matching approach under different conditions. We structure our work by comparing the inference performance of parametric and non-parametric inference methods as described in Figure 2.

**Figure 2:**
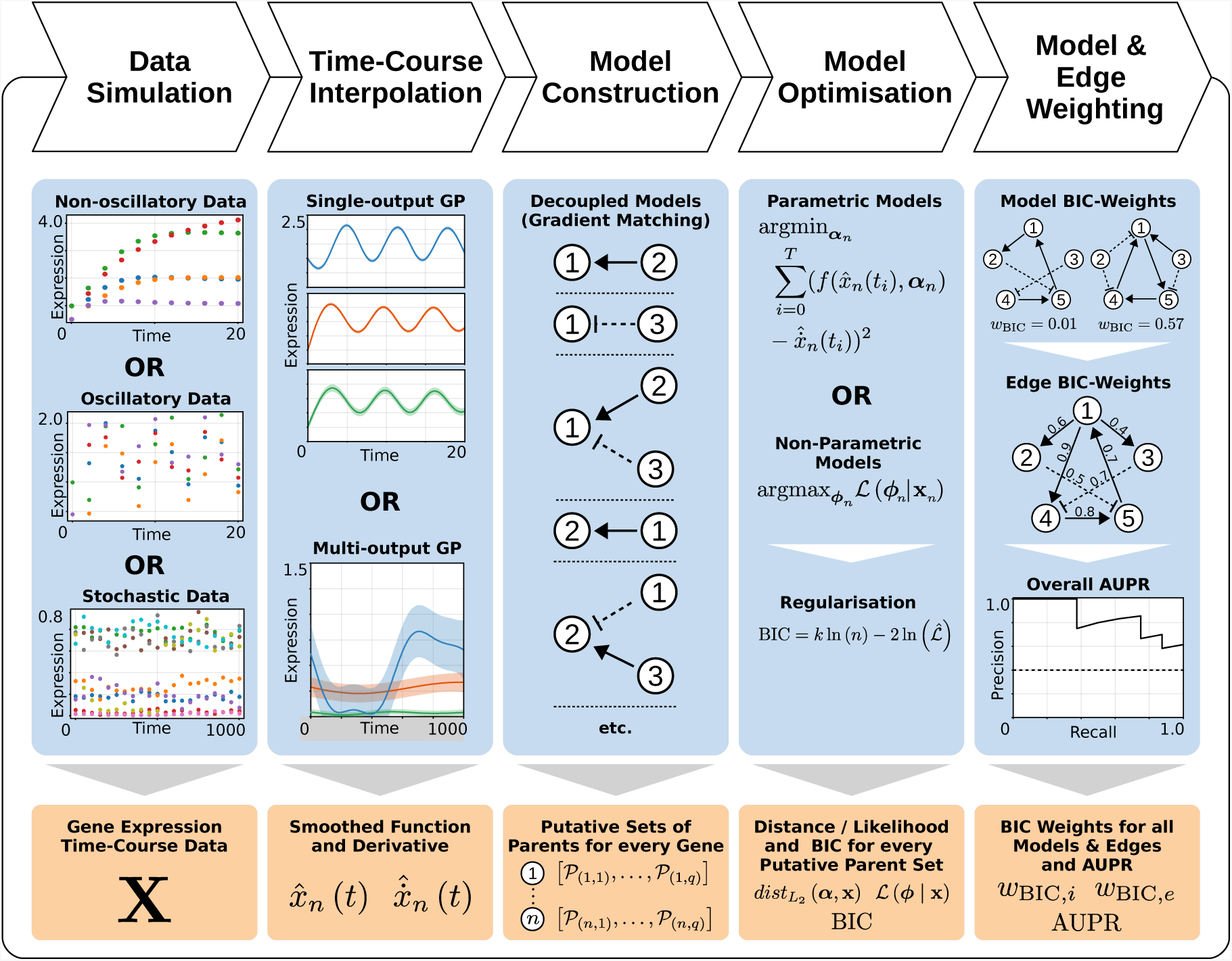
Pipeline outline schematic. This figure illustrates the five main steps in the network inference pipeline developed in this project. All numbers and schematics are shown purely for illustration and do not reflect actual results. Abbreviations used here are: **GP** – Gaussian Process; **BIC** – Bayesian Information Criterion; **AUPR** – Area under the precision-recall curve.

## 2 Methods

This section outlines the different approaches taken to reconstruct GRN. Details on the software and algorithms employed can be found in appendix D.

### 2.1 Gene Expression Data

To compare different network inference approaches and settings, we simulate deterministic gene expression data from a relative small 5-gene regulatory network. We then repeat the analysis using more realistic stochastically simulated data generated from a 10-gene regulatory network in *Saccharomyces cerevisiae*.

#### 2.1.1 Deterministic ODE Model Simulation

We use deterministically simulated gene expression data based on the *in vivo* benchmarking of reverse-engineering and modelling approaches (IRMA) network [19]. The IRMA network is a quasi-isolated synthetic five-gene network, constructed in *Saccharomyces cerevisiae* (Figure 3A). We refer to this dataset as ‘non-oscillatory data’.

To ensure comparability to previous work with this model [11, 18], we use the same model parameters and also create a second subset with one edge removed (Figure 3B) and regulatory interactions modelled as previously [11, 18, 20]. We refer to this dataset as ‘oscillatory data’. For completeness we provide the structure of the ODE systems as well as the parameters and settings used for simulation once again in appendix A.1.

#### 2.1.2 *In silico* Gene Expression Data

In order to evaluate the performance of different inference methods with more realistic stochastically simulated gene expression data, we use `GeneNetWaver` [21] to generate realistic gene expression profiles from an *in silico* ten gene network (Figure 3C) from *Saccharomyces cerevisiae* (as previously used in the DREAM3 and DREAM5 challenge [22]). The dataset we used is referred to as `InSilicoSize10-Yeast1 dream4` in `GeneNetWeaver`. We obtain data for the same 20 time points for every gene.

**Figure 3:**
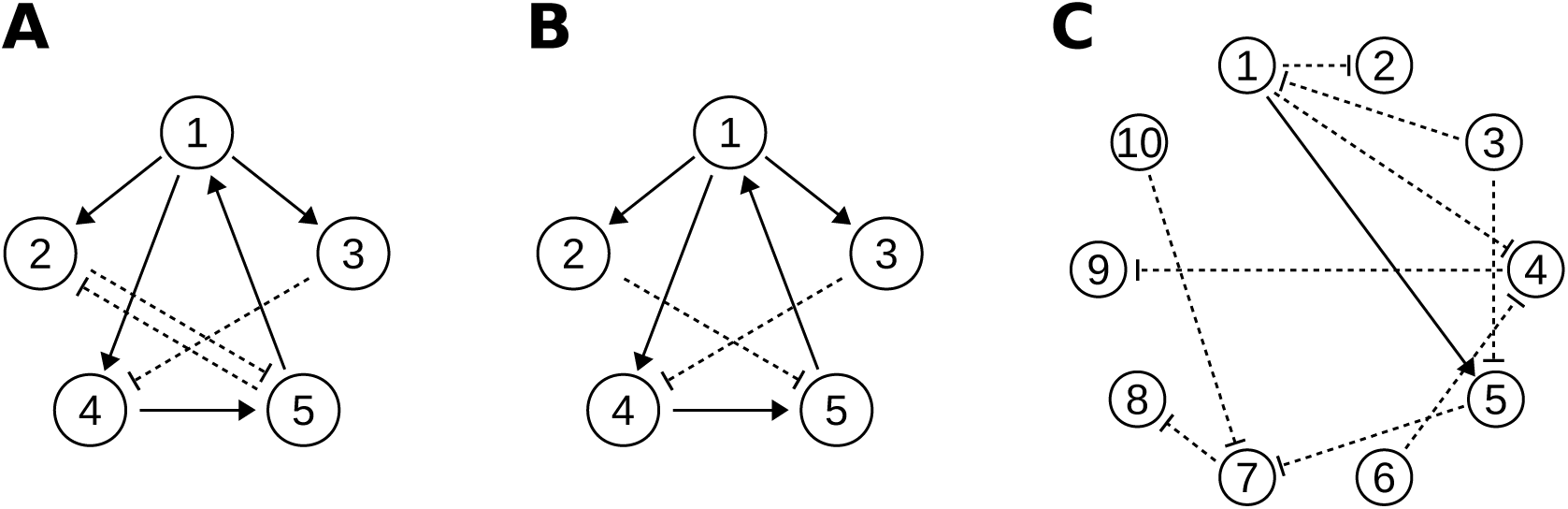
Schematics of gene regulatory networks used in this work. **A** Five-gene network with eight interactions used to simulate the ‘non-oscillatory noise-free data’. **B** Five-gene network with seven interactions used to simulate the ‘oscillatory noise-free data’. **C** Ten-gene network with ten interactions used by `GeneNetWeaver` to simulate the ‘realistic *in silico* data’.

`GeneNetWaver` [21] simulates realistic noisy gene expression data by introducing process noise (through stochastic differential equations) as well as observational noise to the underlying gene expression profiles.

### 2.2 Data Smoothing with Gaussian Processes

For smoothing and interpolation of the potentially noisy gene expression data, we use Gaussian process (GP) regression. This also allows us to obtain the rate of change in the expression via the GP derivative, which is analytically obtainable. In this section we only provide a very brief introduction to the theoretical foundations of GPs and mainly focus on outlining our choices and settings used in the GP framework. For more detail we refer to [23, 24, 25].

#### 2.2.1 Gaussian Process Regression

A GP is defined by a mean *m* and covariance function *k*, so that we can write 
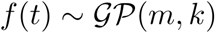
for any suitable function *f*. Any finite collection of values from *f*(*t*) are hence distributed according to a multivariate Gaussian distribution and so we can write 
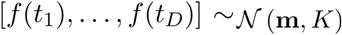
. **m** describes the vector of *n* mean values and *K* = *k* (*t*, *t*′) is the covariance matrix, where the value of each element is defined by the GP covariance function.

We use a zero mean function and employ the common squared exponential covariance function [23], which defines the covariance between two observations at time points *t* and *t*′ as

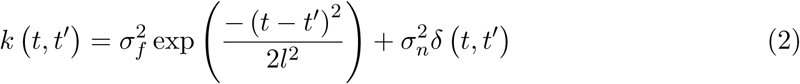

with 
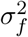
 controlling the variance (‘amplitude’) of the the GP, and the length-scale *l* controlling how many data points around the current one are taken into account when fitting the GP. 
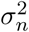
 denotes the variance of the observational noise and we can write 
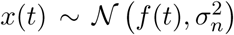
. *δ* (*t*, *t*′) denotes the Kronecker delta function, which is 1 if *t* = *t*′ and zero otherwise. We optimise the hyperparameters *ϕ* = {σ_*f*_,σ_*n*_,*l*} by maximising,

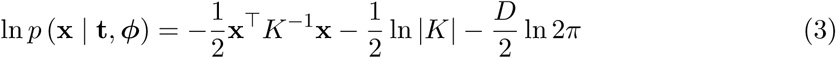

where *K* corresponds to the covariance matrix and *D* denotes the number of observations in vector x. **t**, x ∈ **R**^D^.

We obtain predictions x_*_ at time points 
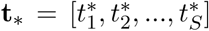
 from the GP model, since the joint (prior) probability distribution of the training output x and testing output x_*_ is again multivariate Gaussian,

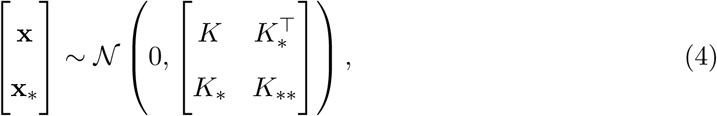

where

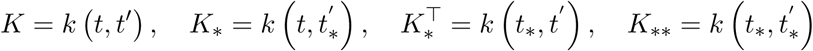

The posterior distribution of the output at t_*_ can be calculated as,

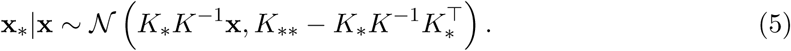

#### 2.2.2 Gaussian Process Derivatives

We can also directly obtain the derivative of the GP, representing the rate of change in mRNA concentration 
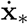
, as the derivative of a GP is again a GP [24, 26],

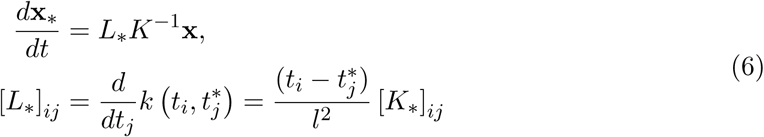

The derivatives obtained here will also be used for gradient matching inference algorithm to be discussed next.

#### 2.2.3 Multiple Output Gaussian Processes

Standard GP regression allows us to make predictions on the expression level of a single gene. To improve the GP fitting to multiple genes, intrinsic coregionalisation for multi-output GP regression [27] ia employed. This is a form of a multiple output GP [28] which takes into account correlation between the expression of all genes in the network through a correlated noise process. Considering a system with *N* outputs, the overall covariance (or kernel) matrix *K* of the multi-output GP takes the form,

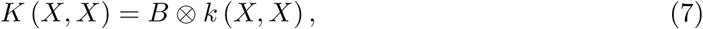

where *B* ∈ **R**^*N*×*N*^ is the coregionalisation matrix, 
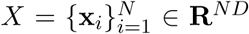
is the input vector that contains observations for all the *N* outputs, and ⊗ denotes the Kronecker product. If *B* = *I*_*N*_, then all outputs are uncorrelated. The hyperparameters in the covariance function *k*(*X*, *X*) and *B* can be estimated jointly via the eigen-decomposition of the matrix *B* and maximum likelihood estimation [29].

We obtain the smoothed mRNA concentration values from the mean function of the GP. We can approximate the derivative at each point numerically,

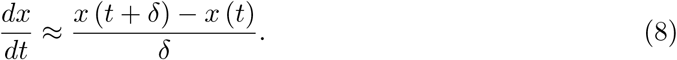

Here we use *δ* = 10^−4^.

### 2.3 Model Construction and Optimisation through Gradient Matching

We use a gradient-matching parameter optimisation approach to evaluate the goodness of fit of our model to the data [11, 14, 18]. Instead of solving the ODE systems, we directly compute the gradient of the gene expression data using GP regression (see Section 2.2) and then optimise the parameters of the ODE system.

As gradient matching can be carried out for each equation of the ODE system independently, the number of possible network topologies we have to consider reduces drastically. For the five gene network with two alternative interaction types and no self-interactions, we only have to consider 
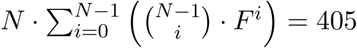
 topologies, given the decoupled system (opposed to *N* = 3.5 · 10^9^ fully coupled models). We can further limit the number of topologies by restricting the number of maximum parents per gene (e.g. the maximum in-degree of every gene in the network). A reasonable assumption would be a maximum of *M* = 2 parents per gene, which would further reduce the space of candidate topologies to 
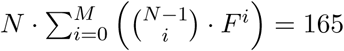
.

### 2.3.1 ODE Models

As during data simulation (Section 2.1.1) we use two different approaches to model activation and repression during network inference. The parameters and constraints used for model optimisation are provided in the Appendix A.2.

For the *n*^*th*^ ODE we minimize the *L*_2_ (squared) distance between the constructed parametric function 
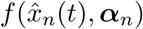
 (with parameter vector *α_n_*) and the associated derivative calculated from the GP regression 
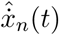
 for all *S* time points [*t*_0_,…, *t_S_*]:

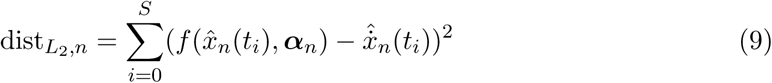

#### 2.3.2 Non-Parametric Models

We also consider a fully non-parametric, GP-based gradient matching inference method adapted from [14]. This is particularly useful when the detailed reaction kinetics (i.e. ODEs) are unknown and when we are more interested to infer the network interactions instead of the kinetics or reaction types (i.e. activation or repression). Similar to the decoupled ODE system described in the previous section, gradient matching approach can also be integrated with non-parametric GP regression. This allows for treating each gene *n* conditionally independent of all other genes given its parents 
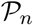
. We model each gene using the relationship:

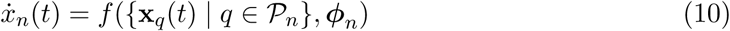

Where 
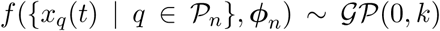
 is a single-output-multiple-input GP with *ϕ_n_* denoting the vector of hyper-parameters for the squared exponential covariance function *k* (*t*, *t*′) (equation 2) for gene *n*. The derivative of the *n*^*th*^ gene expression 
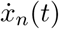
 can again be obtained from the derivative GP process. Optimisation of each putative GP model is via optimising the hyper-parameters of the covariance function by maximizing the likelihood function.

As this is a purely data-driven approach, basal transcription and degradation are not treated separately as in the ODE approach. Because the degradation of mRNA is usually modelled as a first order reaction, we include gene self-interaction in every putative network. This does not affect the total number of candidate topologies. Furthermore, as this approach is unable to distinguish alternative regulatory types (activation or repression) between genes so that the number of possible network topologies is reduced to N 
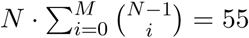
 (with *M* = 2 and *N* = 5).

### 2.4 Model Selection and Edge Weighting

Following model optimisation, we obtain the final distance or likelihood of each gene with respect to their possible parents. For the ODE-based inference approach (Section 2.3.1) we have,

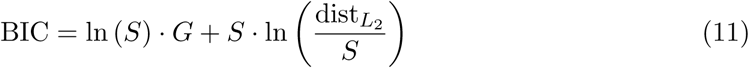

where *S* denotes the number of data points (sample size), *G* the number of free parameters and *dist*_*L*_2__ the *L*_2_ distance defined in equation 9. Alternatively, for the non-parametric inference approach (section 2.3.2) we obtain,

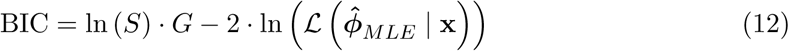

*S* and *G* are defined as before and 
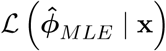
 denotes the maximum likelihood of the model with optimised hyperparameters 
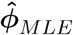
 given gene expression data x. We use the BIC for weighting candidate models rather than the commonly used Akaike information criterion (AIC), as it is asymptotically valid for large sample sizes [30] whereas AIC tends to prefer overly complicated models in this case.

We then calculate the Schwarz weight [31] for each model *w_i_* (BIC) in the set of models *j*,

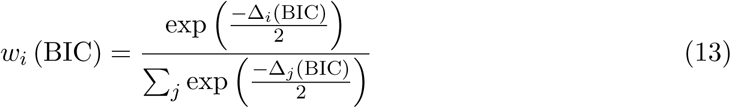

such that Σ*_i_ w_i_* (BIC) = 1. Δ_*i*_ (BIC) = BIC_*i*_ − BIC_*min*_ denotes the difference between the BIC of model *i* (BIC_*i*_) and the lowest BIC across all models considered (BIC_*min*_).

Once we have weighted all models across all genes in the network, we can calculate the weight *w_e_* associated with every edge e in the GRN. This is done for each edge by summing the Schwarz weight of every model that contains the edge in question,

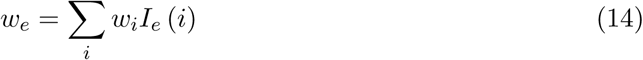

where *I*_*e*_ (*i*) denotes the indicator function which is 1 if edge *e* is present in model *i* and 0 otherwise.

### 2.5 Performance Evaluation

To evaluate the overall performance of the GRN inference, we use the BIC weights of every edge in the network to calculate the Area Under the Precision-Recall (AUPR) curve [32]. The detailed explanations and definitions of this AUPR approach are provided in the Appendix A.3.

Considering the sparsity of large GRN, we use the AUPR instead of the Area Under the Receiver Operating Characteristic (AUROC) curve [33] to evaluate performance.

## 3 Results

### 3.1 Deterministically Simulated Gene Expression Data

For the deterministically simulated gene expression data (see Section 2.1.1), we compare three main approaches to network inference (Table 1: ‘Inference method’). All three methods are combined with gradient matching as described in Section 2.3. For each inference approach, we evaluate a range of different settings (Table 1) using the AUPR (as explained in Section 2.5). For the detailed model and parameter settings, please see Appendices A.1 and A.2. All data presented in this section represent the mean of five independent repeats.

**Table 1:** Employed settings for different network inference approaches.

| Parameter | Settings |
| --- | --- |
| Inference method | Gaussian Process only (non-parametric),<br>ODE constrained parameters,<br>ODE unconstrained parameters |
| Input data | Non-oscillatory data (deterministic, 5 genes, 8 interactions),<br>Oscillatory data (deterministic, 5 genes, 7 interactions),<br>Realistic *in silico* data (stochastic, 10 genes, 10 interaction) |
| Data interpolation | Independent single-output GPs, Multiple-output GP |
| Number of datapoints | 21, 41 |
| Max. num. of parents | 2, 3 |
| Fixed GP length-scale<br>(real. *in silico* data only) | 50, 100, 150, 200 |

**Figure 4:**
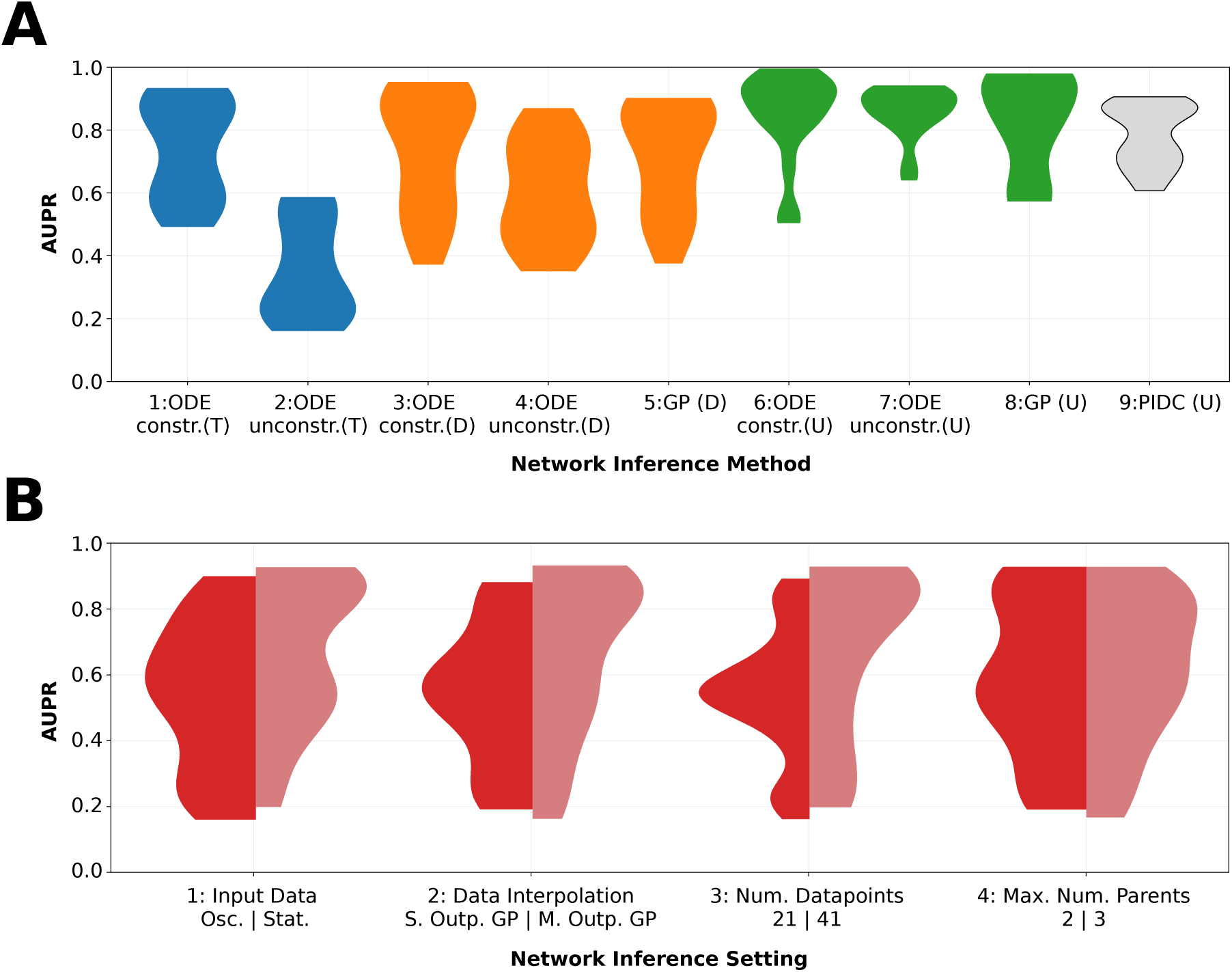
Performance comparison of network inference approaches using noise-free data. **A:** This subfigure displays the distribution of obtained performance (**AUPR**) for the three different classes of network inference methods, over all model settings listed in Table 1. There are four different network inference aims shown in four different shades. The blue distributions relate to the performance of the constrained and unconstrained ODE methods at inferring a directed GRN including information about interaction types (activation/repression) (**T**). The orange distributions depict the performance of the two ODE-based methods and the GP-based method at predicting a directed GRN without type information (**D**). The green distributions show the performance of the same three methods at inferring an undirected GRN (**U**). The performance of a recently developed algorithm [8] based on partial information decomposition for the same settings and data is shown as the last distribution in grey (“PIDC”). *Baseline (random) performance is 0.2 for plots 1-2, 0.4 for plots 3-5, 0.7 for plots 6-9*. **B:** This subfigure shows the impact of different settings choices on network inference performance. Summing the two halves of each of the four asymmetric distributions in the figure gives rise to the same distribution of model performance (constituted by the three approaches discussed earlier - represented distributions 1, 2 and 5 in A).

#### 3.1.1 Comparing Parametric and Non-Parametric Inference

Figure 4A contrasts the performance the three inference approaches across all settings and for three different inference aims, respectively. Only the parametric ODE-based methods allow for distinction between activating and repressing regulatory interactions between genes. From Figure 4A, we can however clearly see that this type of inference is successful only if the detailed kinetic information about the GRN is available prior to inference: the unconstrained ODE-based modelling of interactions shows a significant drop in performance over the tested settings compared to the constrained approach where basal transcription and degradation rates are known and ODE parameter ranges can be constrained *a priori* (see Table 2 in Appendix A.1 for parameters).

If we are only interested in the directionality of interactions and not their specific type, the three orange distributions in Figure 4A show that constraining the parameters of the ODE-based approach (and assuming known basal transcription and degradation rate) is no longer important for achieving good inference performance. The GP-based approach achieves on average higher performance on the simulated datasets used here. This is surprising, since gene interactions used in generating the data are of the same functional form assumed in the ODE inference.

The same trend (with slightly higher overall performance) can be seen when we are only predicting undirected edges. Interestingly, despite higher overall performance, constraining the ODE parameters can lead to worse performance under certain inference settings for this task (compare plot 6 and 7 in Figure 4A). All three approaches generally perform better on this simple noise-free five-gene networks than the PIDC approach [8].

Below, we analyse the impact of individual factors on inference quality, such as the data interpolation method, on the overall performance of the discussed methods.

#### 3.1.2 Input Data

The distributions separated by the two input data types (plot 1, Figure 4B) show a slight performance increase for the non-oscillatory dataset over the oscillatory one. This should be taken into account during experimental design in order to produce data which bears maximum information about the underlying regulatory network [34].

#### 3.1.3 Data Interpolation

Despite the deterministic nature of the data we use for evaluation in this section, we find a pronounced difference in performance depending on the method used for interpolating the input data. By taking into account the correlation between the different gene expression time-courses, interpolation with a multiple output GP is able to achieve significantly better results compared to using independent GPs.

When interpolating oscillatory data using single output GPs, we observe that for low number of data points, the GP hyperparameters are optimised so that the oscillatory behaviour is no longer traced by the GP mean, but rather interpreted as noise (figure 14A in Appendix C). This was also observed in previous work [11]. As shown in Figure 14B (appendix C)this problem can be overcome by using multiple output GP regression, where the oscillatory behaviour correctly traced because trajectories of all genes are taken into account when optimising hyperparameters [35, 36].

#### 3.1.4 Number of Data Points

Plot 3 of Figure 4B demonstrates increased performance as more time points are used. While this is unsurprising for noise-free data, we will re-evaluate this observation for stochastic data in Section 3.2.2.

#### 3.1.5 Maximum Number of Parents Considered

In figure 4B we can see that the maximum number of parents considered per gene does not markedly affect performance. From this we conclude that for deterministic data, these network inference methods may be robust even for large putative network spaces. The main reason for this robustness is the weighting of the models using the BIC as this metric penalises model complexity explicitly.

### 3.2 Noisy *in silico* Gene Expression Data

Gene expression is a stochastic process and we apply the same inference procedures to stochastically simulated *in silico* gene expression data (but for 10 instead of 5 genes; see Section 2.1.2 for details).

#### 3.2.1 Comparing Parametric and Non-Parametric Inference

The most notable difference between the results for the noise-free and noisy gene expression data is the absolute decline in performance, which is not unexpected. Despite this difference, we nevertheless observe similar trends as for the noise-free data. The unconstrained ODE-based modelling (plot 2, Figure 5A) again provides comparable performing result to the non-parametric GP-only modelling approach (plot 3, Figure 5A) when interaction types are not of interest.

When trying to infer only the existence of (undirected) edges between genes, we observe that the unconstrained ODE-based model performs slightly better than the GP-based approach; and both approaches perform better than PIDC.

The pronounced narrowing of distributions towards higher AUPR across different approaches indicates that unlike inference based on noise-free data, both ODE and GP-based methods only produce meaningful results (i.e. significantly better than random performance) for a very narrow range of scenarios.

#### 3.2.2 Model Settings

Contrasting the performance for noise-less and noisy data shows not just lower absolute performance for each method for noisy data, but also different trends of their behaviour (Figure 5).

Interestingly, we can see from plot 1 of Figure 5B that in case of stochastic data, all well-performing inference approaches use single output GP interpolation of the data. This could be explained by the large number of free parameters in multiple output GP optimisation. For a ten-gene network, moving from ten independent single output GPs to one 10-output GP means solving a 32-parameter optimisation problem (31 for fixed length-scale) in contrast to solving ten 3-parameter problems. As finding the optimal solution in such a high-dimensional parameter space is extremely difficult, this may be the leading cause for this observation. We further substantiated this by interpolating gene expression data from a smaller GRN using single‐ and multiple output GP regression and comparing network inference results. An additional reason for the reduced performance could be the limitation to a single length-scale hyperparameter for multiple output GP, while single output GPs can have a different length-scale and variance for every gene they fit. This allows for more flexibility during interpolation. Multiple-output GP methods which allow for varying length-scales are available [37], however, but this further increases the number of free hyperparameters to be optimised.

**Figure 5:**
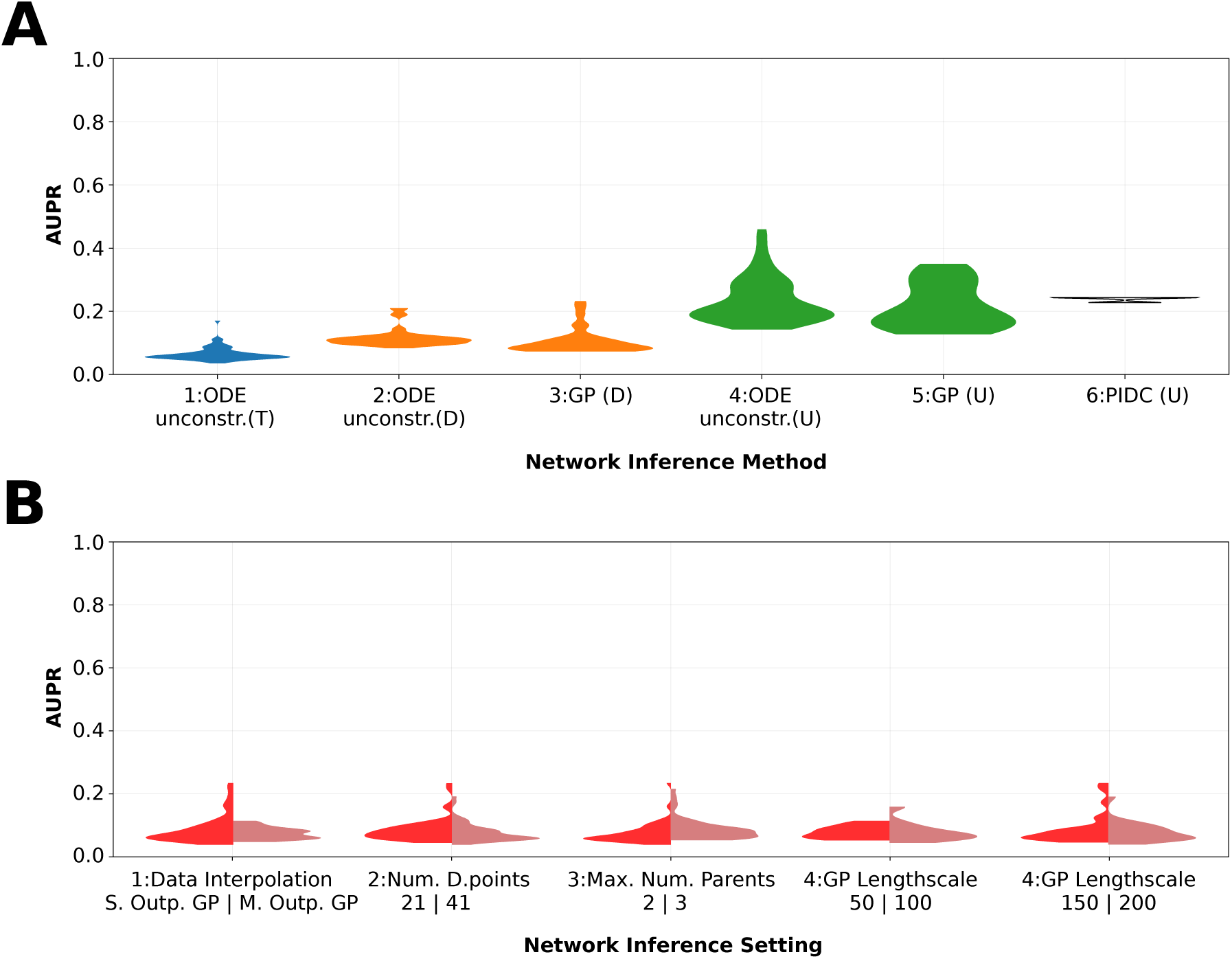
Performance comparison of network inference approaches using realistic simulated *in silico* data. **A:** This subfigure displays the distribution of obtained performance (**AUPR**) for the three different classes of network inference methods, over all model settings listed in Table 1. There are four different network inference aims shown in four different shades. The blue distribution relates to the performance of the unconstrained **ODE** method at inferring a directed **GRN** including information about interaction types (activation/repression) **(T)**. The orange distributions depict the performance of the **ODE**-based method and the **GP**-based method at predicting a directed GRN without type information **(D)**. The green distributions show the performance of the same three methods at inferring an undirected GRN **(U)**. The performance of a recently developed algorithm [8] based on partial information decomposition for the same settings and data is shown as the last distribution in grey **(“PIDC”)**. Baseline (random) performance is 0.06 for plot 1, 0.1 for plots 2-3, 0.2 for plots 4-6. **B:** This subfigure shows the impact of different settings choices on network inference performance. Summing the two halves of each of the first three asymmetric distributions in the figure gives rise to the same distribution of model performance (constituted by the two main approaches discussed earlier (GP and ODE) - represented by distributions 1 and 5 in Figure 5A). The same is true for the sum of the last two distributions in the figure.

We also see from plot 2 of Figure 5B that increasing the number of data points taken from the interpolated data no longer improves performance. While this might seem counter-intuitive at first, the inability of the GP to interpolate the true underlying gene expression dynamics renders the benefit of more data points futile; it appears that GPs can overfit the noise in the data (unless the GP hyperparameters are specifically constrained); using fewer time points can partially compensate for such overfitting. On closer inspection we find that this effect is particularly pronounced for the derivatives obtained from the GPs that play a major role in the inference.

Again changing the maximum number of parents allowed for a gene appears to have no effect (plot 3, Figure 5B). The rightmost two plots of Figure 5B show clear evidence for the importance of the right choice of length-scale during data interpolation (only at a length-scale of 150 can an inference performance of AUPR > 0.2 be achieved for this example).

## 4 Discussion

In this work, we compare the performance of different network inference methods, especially parametric and non-parametric gradient matching methods, under different settings and scenarios in order to gain an understanding of the strengths, weaknesses and impact of different modelling choices.

When inferring GRNs from limited and inherently noisy gene expression data, there are usually a large number of potential models that can match the data [18]. By computing weights for each model and consequently each interaction in the network, we are able to obtain useful inferences by pooling over different methods.

We find that the simple non-parametric inference approach achieves slightly lower performance than the unconstrained ODE method despite the absence of mechanistic knowledge about the underlying regulatory processes. It was however shown in previous studies, that a more advanced non-parametric approach which combines Bayesian linear regression and GPs is able to achieve higher performance [14] assuming that some of the parameters are known. In our work, we show that knowledge of such parameters prior to network inference can strongly increase performance and even allows us to infer mechanistic aspects of interactions from data.

When inferring networks from gene expression data, the ability of the GP to reconstruct the underlying time-courses from noisy data is a critical factor. Especially the gradient obtained from the GP for the gradient matching procedure is particularly sensitive to poor fits. In order to alleviate this, previous work [11, 38] has suggested employing adaptive GPs which can improve performance by taking into account the structure of the ODE model (in case of parametric modelling) during GP fitting.

We believe that this approach is still worth pursuing further. Another promising avenue we see for future work is the combination of parametric and non-parametric methods. A possible approach would be to use the computationally cheaper GP-based method to sufficiently narrow the space of possible networks. We could then use ODE-based network inference to confirm interactions as well as obtain mechanistic information for the predicted edges in the GRN. If the space of putative networks is small enough following the non-parametric step, we could even avoid decoupling the network which would further increase inference performance.

## 5 Conclusion

In this work, we have carried out a comprehensive comparison of a range of parametric and non-parametric gradient-matching-based approaches on gene regulatory network inference from gene expression data.

We found that applying parametric ODE-based approaches on deterministic gene expression data showed that mechanistic information (such as the type of interaction) can be recovered during inference if enough knowledge about the network (e.g. parameter ranges) is present. For directed and undirected network inference, parametric ODE method can provide comparable or even better inference performance compared to non-parametric GP-based method, the latter approach however requires little mechanistic or kinetic regulatory information and computationally more efficient, which can be crucial for large-scale network inference problems. When applying to larger network or stochastic data, overall lower inference performance is observed for all methods, while consistent comparable performance between parametric and non-parametric methods is still obtained.

Several promising avenues to improve inference performance emerge from this analysis: in particular there is potential for the use of multiple output Gaussian Processes for data interpolation in cases of small networks. When applying the same methods to more complex stochastic networks these may, however, become less reliable.

A central result has been that Bayesian model averaging has real potential to increase the quality of network inference. We believe that combining the strengths of several existing approaches will ultimately be required to make significant further progress in solving this challenging problem.

## A Appendix: Methodology - Further Details

## A.1 Parameters, Functional Forms and Data for ODE Model Simulation

For simulation of the non-oscillatory data, we model the expression of each gene in the network shown in (Figure 3A) as ODEs of the form (as used in the DREAM7 (Dialogue on Reverse Engineering Assessment and Methods) challenge [39]):

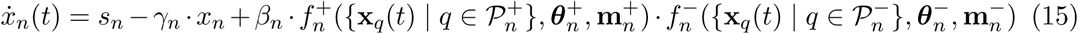

with the following two expressions defining the activating 
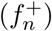
 and repressing 
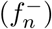
 regulation of gene *n* respectively:

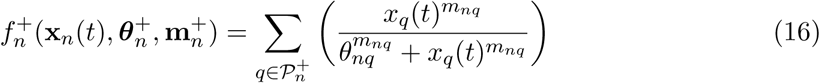

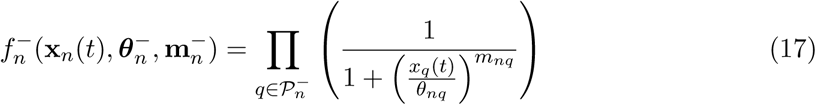

Here, 
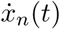
 denotes the rate of change of the mRNA concentration at time *t*, *x*_*n*_(*t*) the mRNA concentration, *s_n_* the basal transcription rate, *γ_n_* the mRNA degradation rate, *β_n_* the strength of gene regulation, all with respect to gene n ∈[1,…,*i*] where *i* denotes the number of genes in the network. 
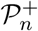
 and 
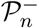
 denote the subsets of parent genes which have an activating or repressing effect on the expression of gene *n* respectively. θ_*nq*_ and *m*_*nq*_ are the commonly used hill parameters for the regulation of gene *n* by its parent *q* [40].

For simulation of the oscillatory data, we model the expression of each gene in the network shown in (Figure 3B) as ODEs of the form (parameter notation as in equation 15):

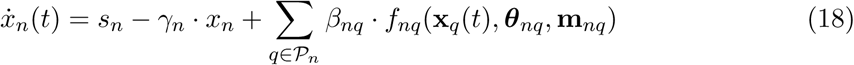

with the following two terms representing the interaction term *f*_*nq*_ of the parent 
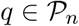
 with the gene *n*.

In case of an activating interaction between the parent gene 
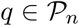
 and gene *n*:

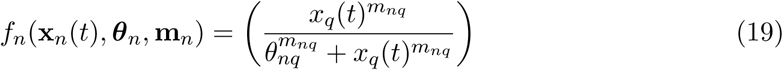

In case of an repressing interaction between the parent gene 
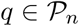
 and gene *n*:

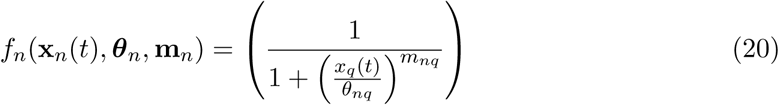

Data simulated from the networks shown in Figure 3 was simulated over the timespan (0,20) with a time-step of either 0.5 or 1.0 for deterministic data and 25 or 50 for stochastic data. Starting concentrations for the five genes [*x*_1_ (0),…,*x*_5_ (0)] were set as [1.0,0.5,1.0,0.5,0.5] in all cases.

The following ODE systems were used for simulation of the oscillatory and non-oscillatory noise-free data:

**Non-oscillatory data**

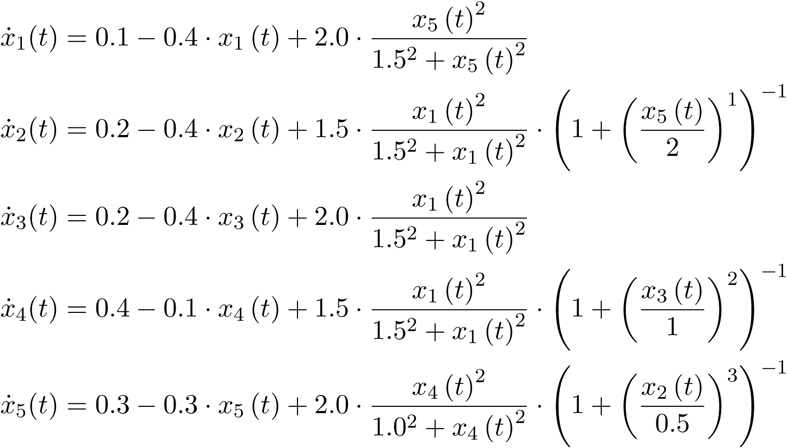

**Oscillatory data**

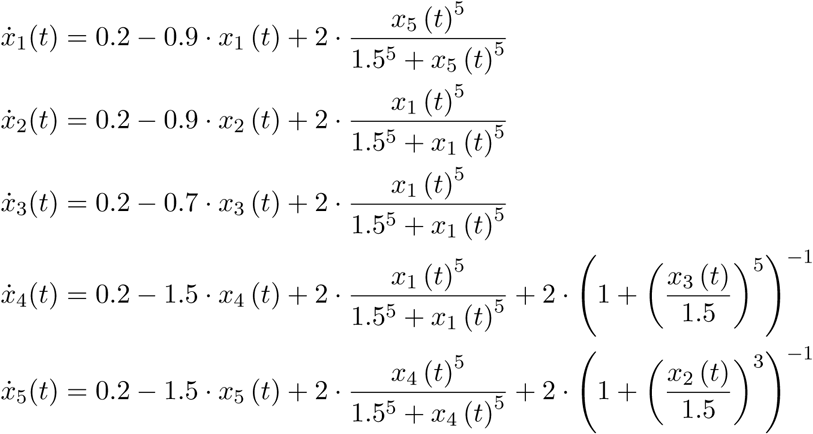

**Raw Gene-Expression Time-Course Data Plots**

**Figure 6:**
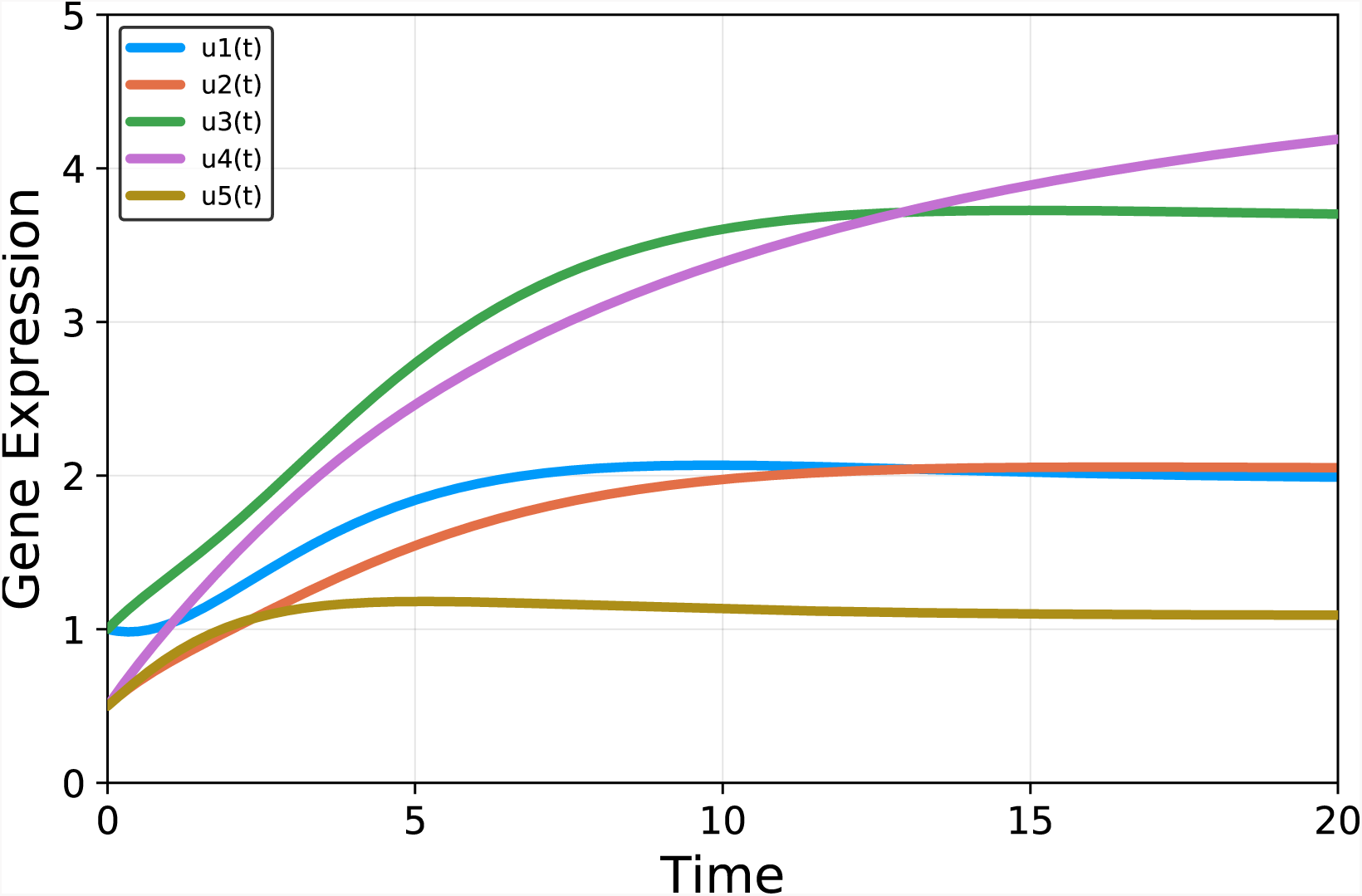
Solution to ODE system used for simulation of the non-oscillatory time-course data.

**Figure 7:**
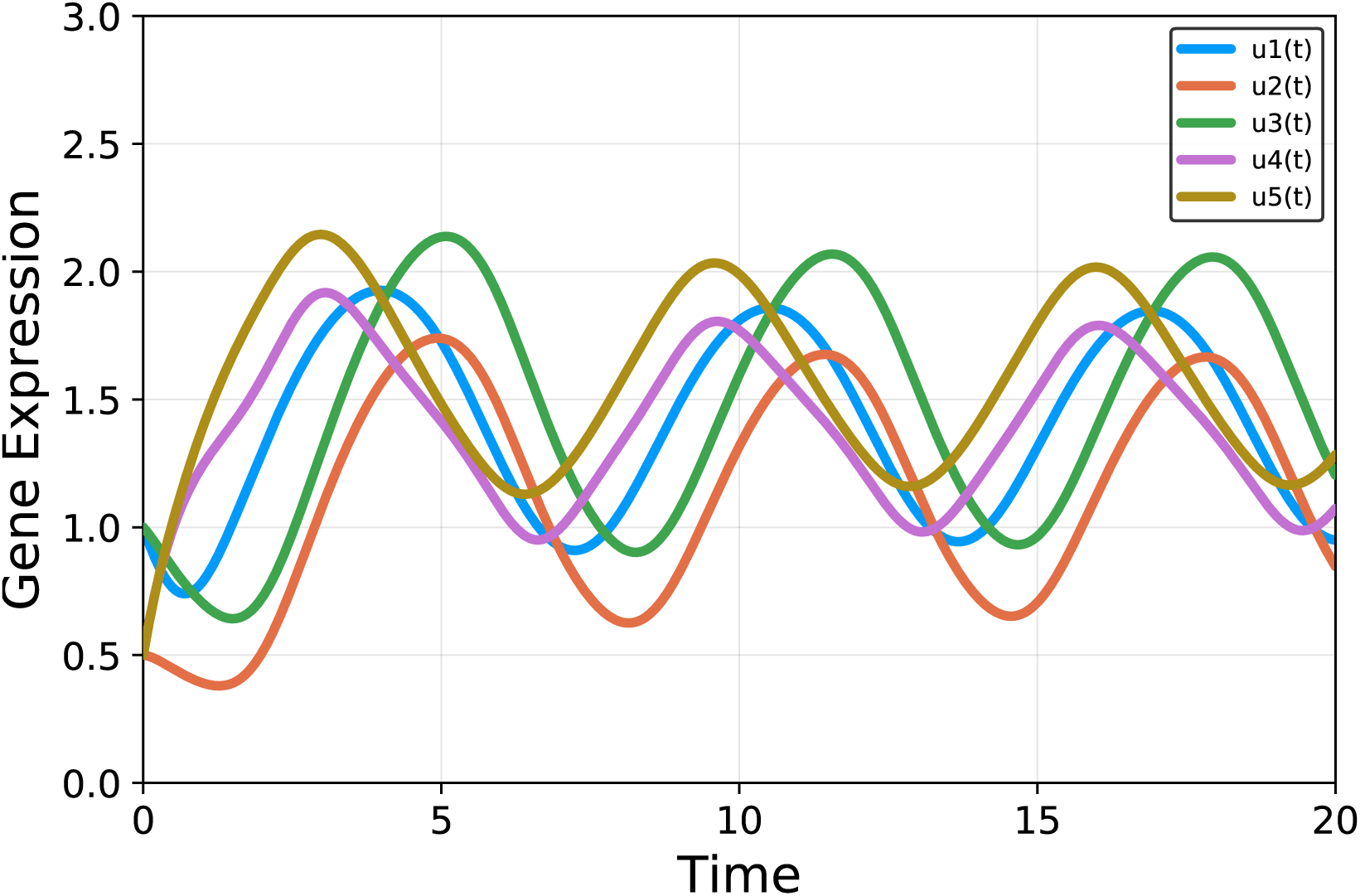
Solution to ODE system used for simulation of the oscillatory time-course data.

**Figure 8:**
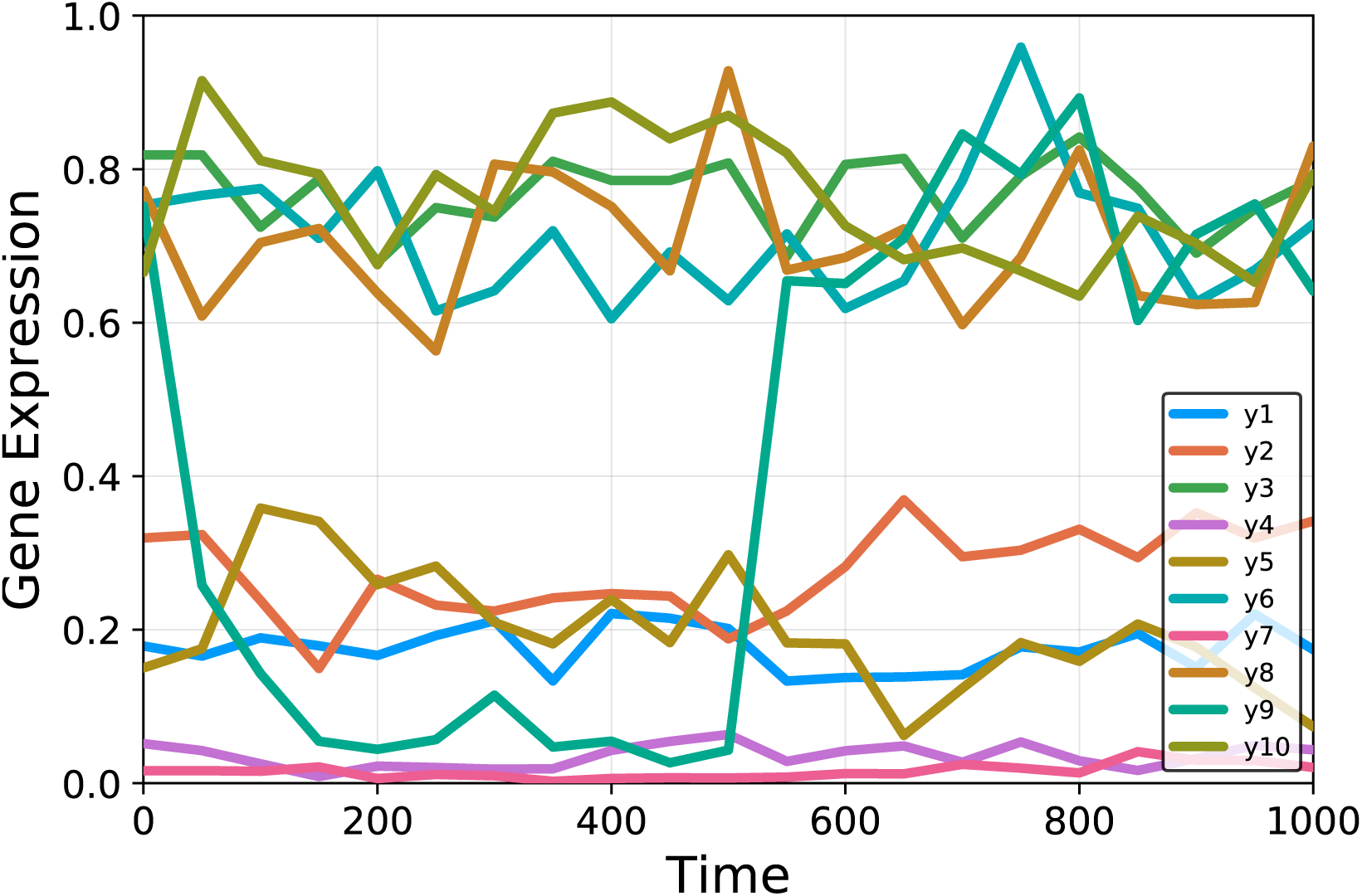
Realistic *in silico* time-course data simulated by `GeneNetWeaver`.

## A.2 Parameters for Model Optimisation

Parameter optimisation of each putative decoupled model was done using either known bounds and starting values for each parameter or completely unconstrained optimisation. For constrained optimisation, basal transcription and degradation rates were assumed to be known, while they were included as free parameters for the unconstrained optimisation.

For the guidelines used to simulate the non-oscillatory data (equations 15, 16, and 17 in Appendix A.1 and Figure 3A), this results in either 3 + 2 · *r* or 1 + 2 · *r* free parameters to be optimised for each decoupled gene, depending on whether basal transcription and degradation rates are known and where *r* is the number of parents for each gene. For the guidelines used to simulate the oscillatory data (equations 18, 19, and 20 in Appendix A.1 and figure 3B), the number of free parameters per decoupled ODE system is 2 + 3 · *r* or 3 · *r* respectively.

Parameter constraints and starting values for the remaining parameters optimised during constrained optimisation are displayed in Table 2. In case of unconstrained ODE optimisation, all free parameters were initialised at 1.0.

For non-parametric network inference, models were optimised using the likelihood (equation 3) as the maximisation objective and starting values 1.0 for all free hyperparameters. Under certain circumstances (as mentioned in the main text) the length-scale hyperparameter of the GP was fixed at 50, 100, 150 or 200 before running the optimisation in order to achieve a satisfactory fit to the data.

**Table 2:** Constraints and and initial parameter values used for constrained optimisation of putative ODE models.

| Data class | Param. | Initial value | Lower bound | Upper b. | True value(s) |
| --- | --- | --- | --- | --- | --- |
| ‘Stat. data’ | $\beta$ | 1.0 | 0.0 | 5.0 | 2.0, 1.5 |
| | $m$ | 2.0 | 0.1 | 5.0 | 1.0, 2.0, 3.0 |
| | $\theta$ | 1.0 | 0.0 | 4.0 | 0.5, 1.0, 1.5, 2.0 |
| ‘Osc. data’ | $\beta$ | 1.0 | 0.5 | 4.0 | 2.0 |
| | $m$ | 1.0 | 0.7 | 5.0 | 5.0 |
| | $\theta$ | 1.0 | 0.2 | 3.0 | 1.5 |

## A.3 Performance Evaluation using Precision-Recall curves

In this section, we explain and define how we evaluated performance of our inference method using precision-recall curves.

To obtain the AUPR, the vector of edge weights was first sorted. Each value in the vector was then iteratively chosen as the cut-off point between a positive and negative prediction (i.e. all larger weights are considered positive predictions and vice versa). It was therefore possible to calculate the precision and recall for each of these thresholds. A graph was plotted using the obtained precision (y-axis) and recall (x-axis) values. The area under the curve arising from this plot is the AUPR (see Figure 9).

The procedure for obtaining the AUROC is analogous, with the only difference that the false positive rate is used instead of the precision.

**Figure 9:**
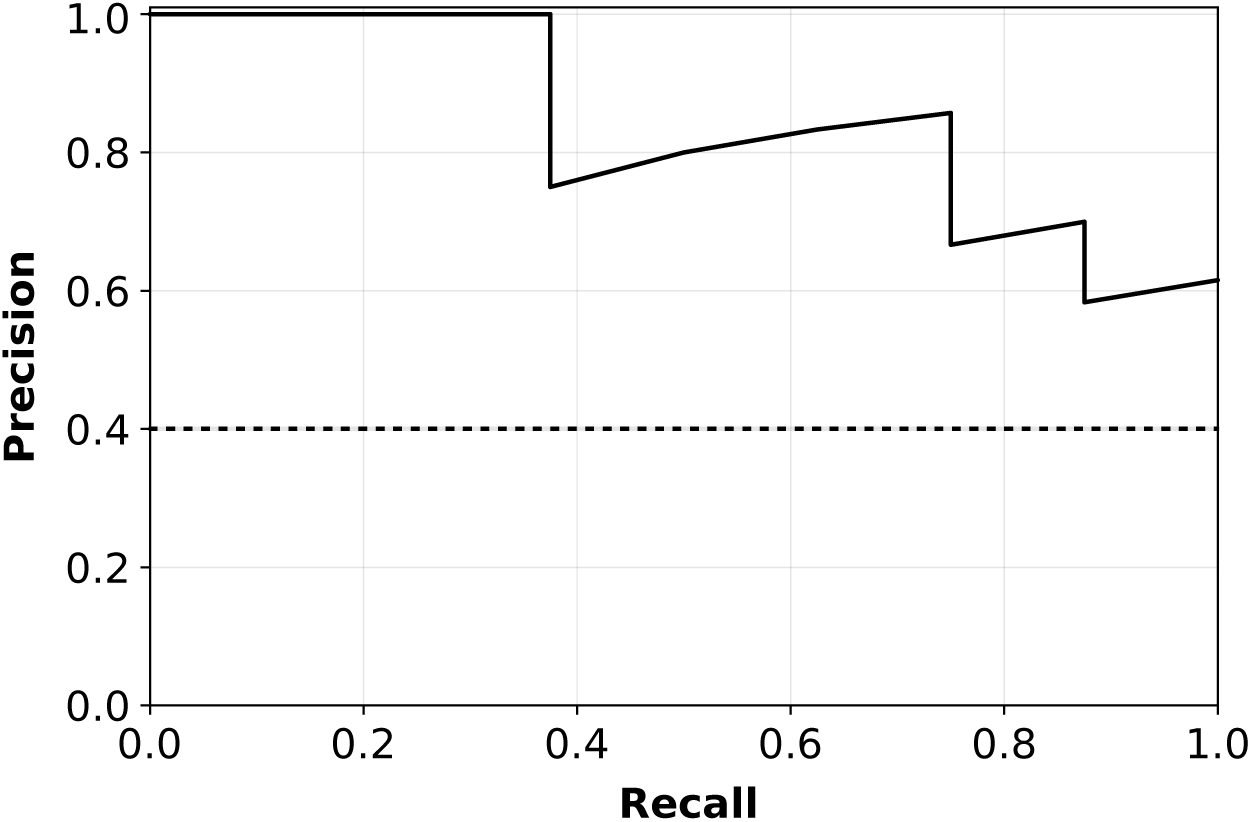
Example Precision-Recall curve. PR curve with AUPR of 0.84 with the baseline (random performance) shown as a dashed line.

**Definitions**

Tp: True positive (a predicted edge which is actually present)

FP: False positive (a predicted edge which is not actually present)

TN: True Negative (an unpredicted edge which is actually not present)

FN: False Negative (an unpredicted edge which is actually present)

*Precision (Positive Predictive Value)* – the proportion of true edges that were predicted correctly:

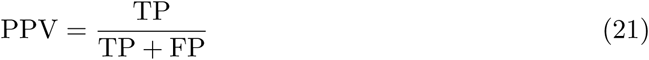

*Recall (True Positive Rate)* – the proportion of predicted edges that are actually present:

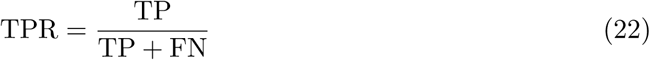

*False Positive Rate* – the proportion of predicted edges that are not actually present:

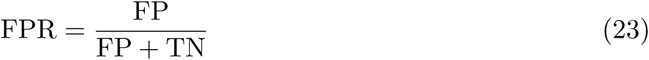

## B Appendix: Supplementary Figures for Settings Evaluation

**Figure 10:**
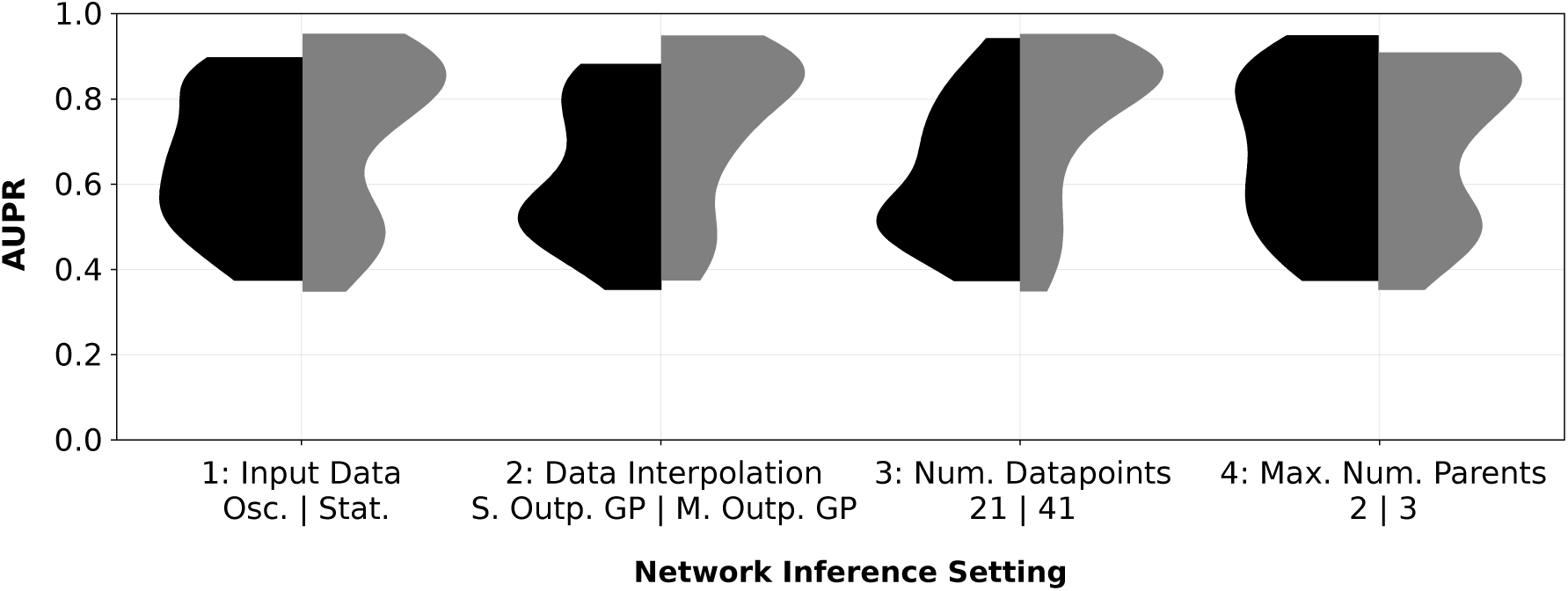
Performance of all evaluated inference approaches by model choices for noise-free data and inferring directed edges. This figure shows the impact of different settings choices on network inference performance. Summing the two halves of each of the four asymmetric distributions in the figure gives rise to the same distribution of model performance (constituted by distributions 1, 2 and 5 in Figure 4).

**Figure 11:**
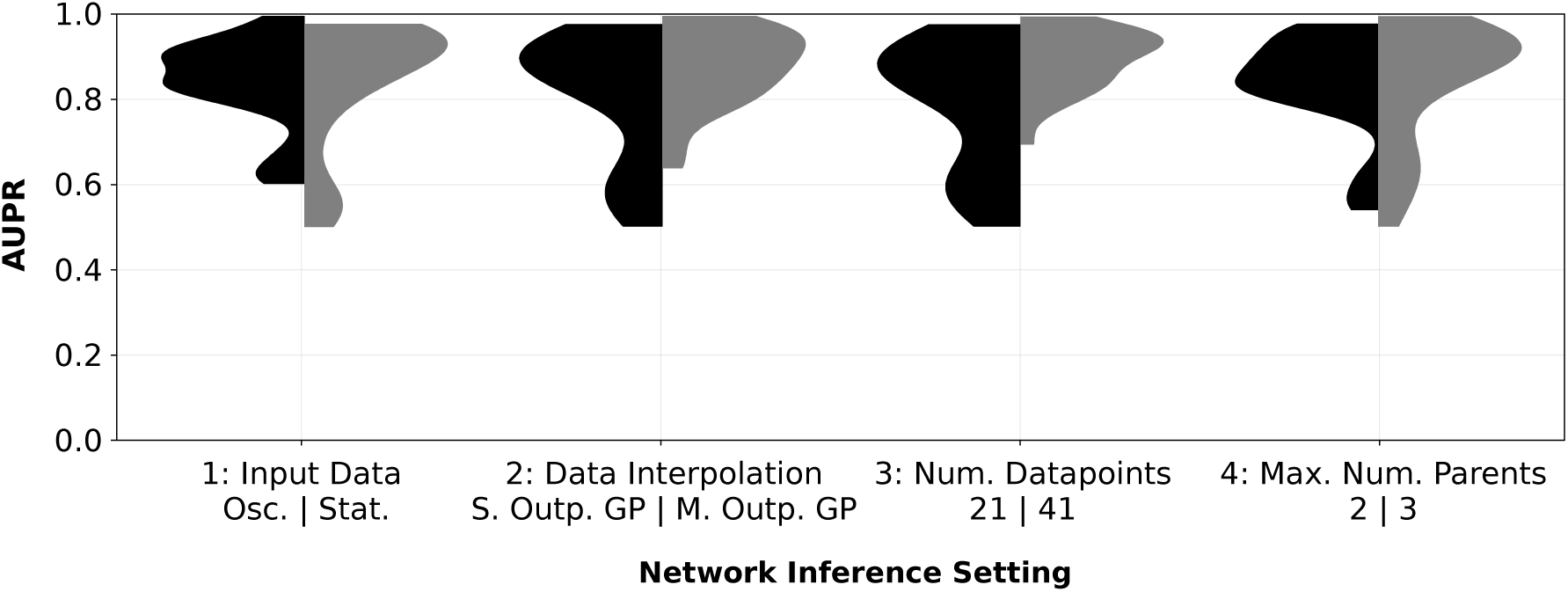
Performance of all evaluated inference approaches by model choices for noise-free data and inferring undirected edges. This figure shows the impact of different settings choices on network inference performance. Summing the two halves of each of the four asymmetric distributions in the figure gives rise to the same distribution of model performance (constituted by distributions 1, 2 and 5 in Figure 4).

**Figure 12:**
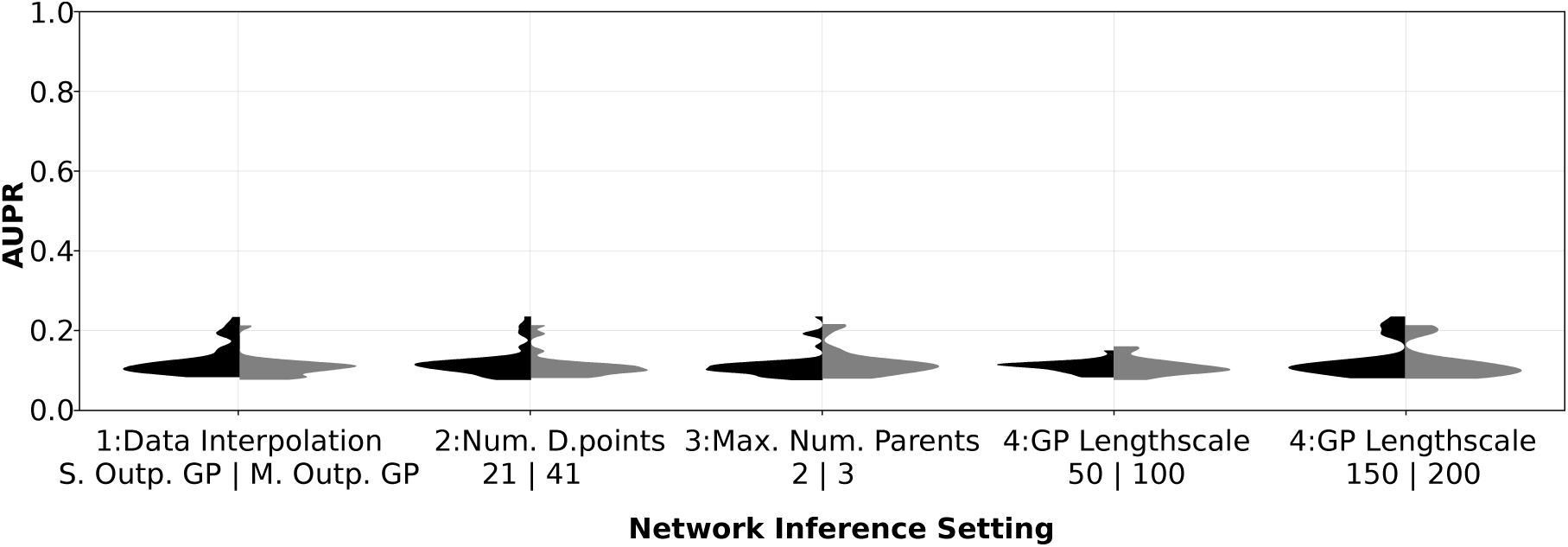
Performance of all evaluated inference approaches by model choices for stochastic data and inferring directed edges. This figure shows the impact of different settings choices on network inference performance. Summing the two halves of each of the first three asymmetric distributions in the figure gives rise to the same distribution of model performance (constituted by distributions 1 and 5 in Figure 5). The same is true for the sum of the last two distributions in the figure (“GP Lengthscale”).

**Figure 13:**
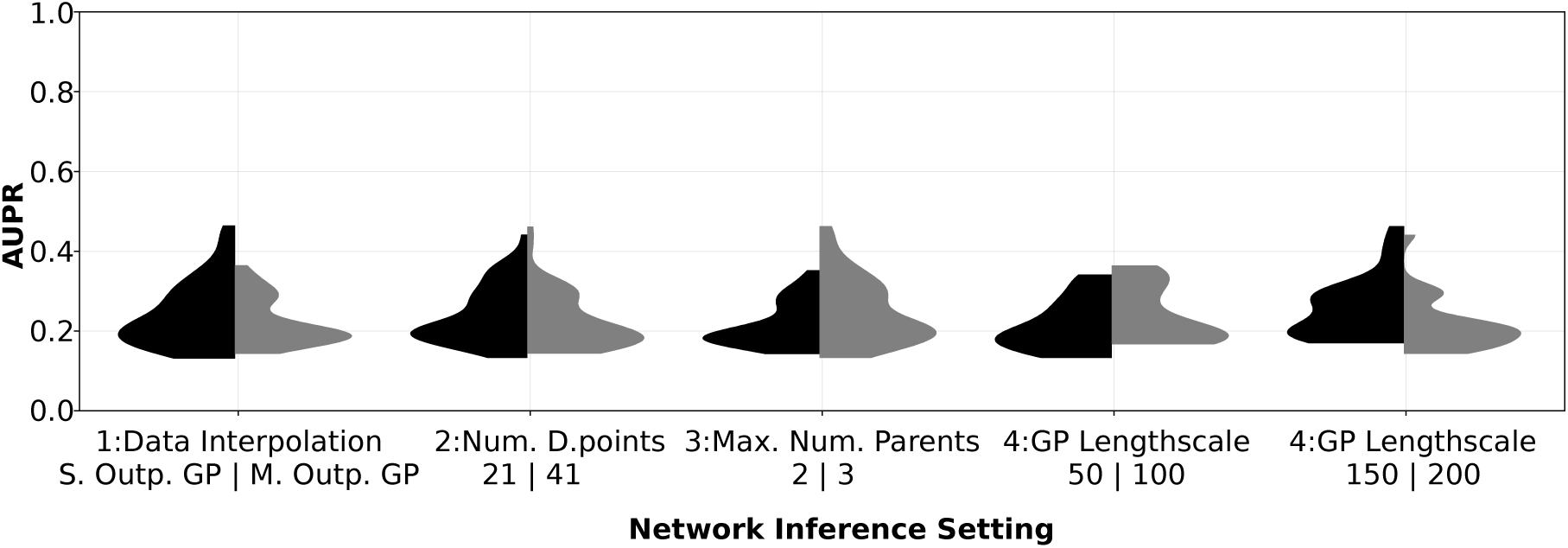
Performance of all evaluated inference approaches by model choices for stochastic data and inferring undirected edges. This figure shows the impact of different settings choices on network inference performance. Summing the two halves of each of the first three asymmetric distributions in the figure gives rise to the same distribution of model performance (constituted by distributions 1 and 5 in Figure 5). The same is true for the sum of the last two distributions in the figure (“GP Lengthscale”).

## C Appendix: Supplementary Figures for fitting GPs

**Figure 14:**
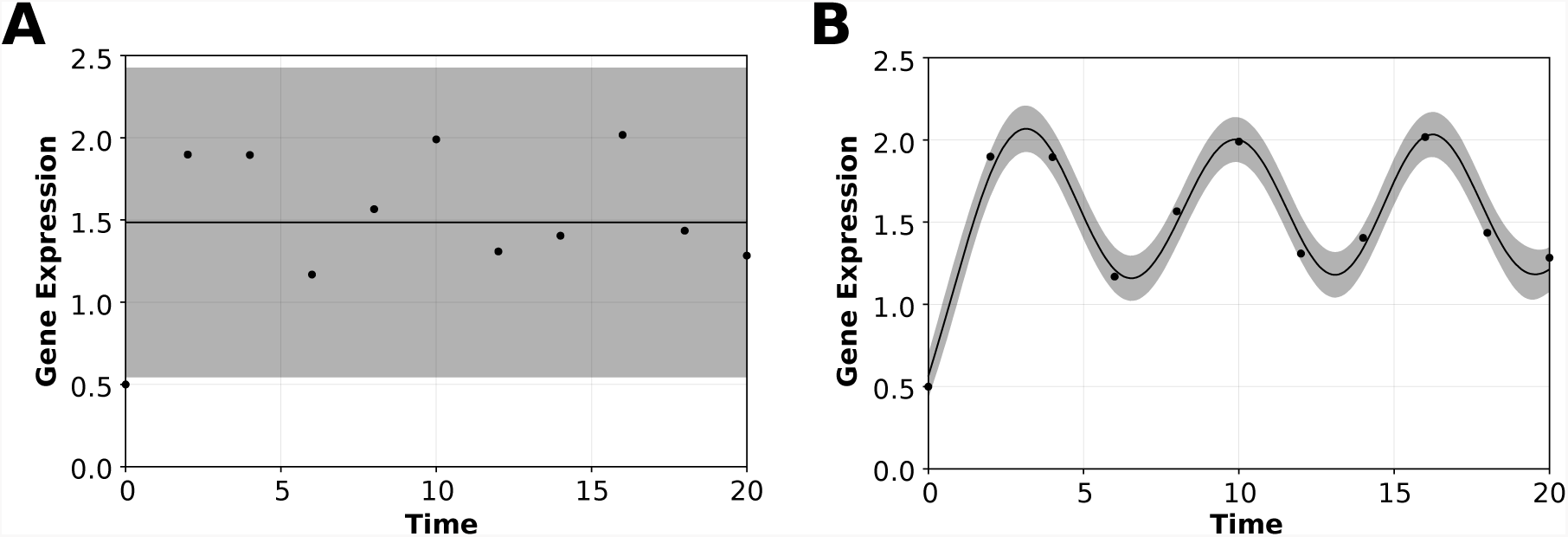
Comparison of single output GP and multiple output GP fits to data. Both plots show time-course data for the expression of gene 5 from the non-oscillatory (noise-free) data indicated by black dots. The black lines refer to the GP mean function with 95% confidence intervals shaded in grey. **A** Data interpolated using a single output GP. **B** Data interpolated using a multiple output GP, which takes into account correlations between the expression time-courses of all genes in the network.

**Figure 15:**
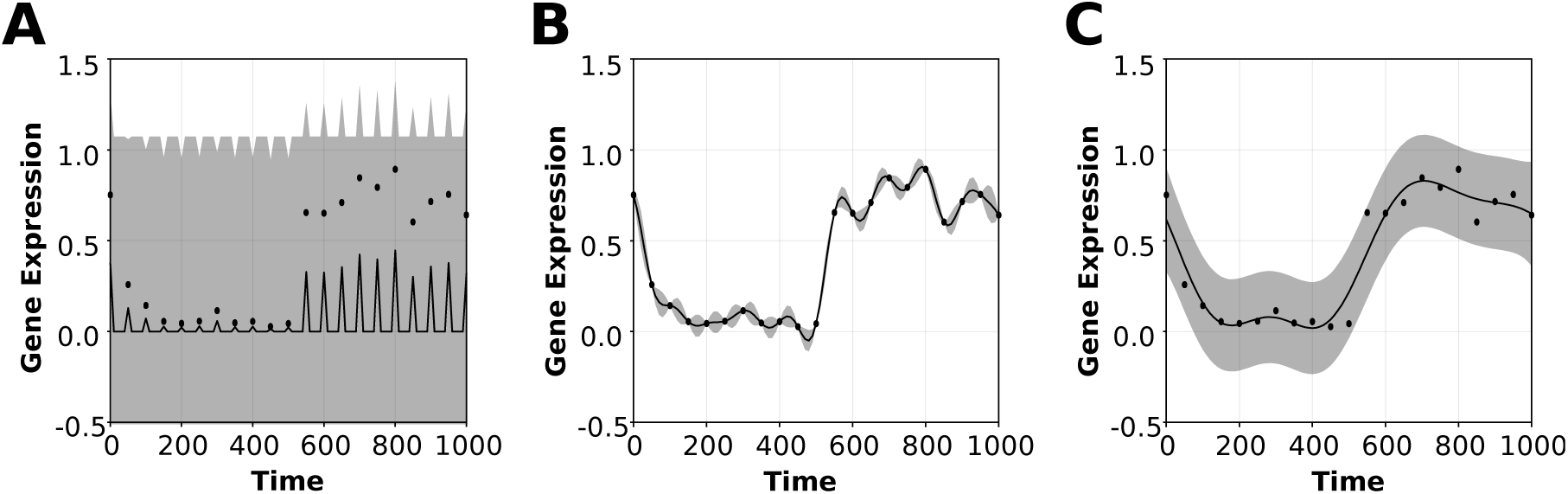
Comparison of GP fit with different length-scale hyperparameters. All three plots show the time-course data for the expression of gene 9 from the GNW dataset indicated by black dots. The black lines refer to the GP mean function with 95% confidence intervals shaded in grey. The three plots show different length-scale hyperparameter settings: **A** unconstrained hyperparameter optimisation, **B** length-scale fixed to 50, **C** length-scale fixed to 150.

## D Appendix: Software Tools

All code for this project was written in the `Julia` programming language, version *0.6* [41].

## D.1 Data Simulation

Simulations of the mRNA expression data described by the ODE systems shown is Section A.1 were carried out using the Runge-Kutta Order 4 (`RK4()`) solver from `Differential Equations.jl`^1^ [42].

## D.2 Gaussian Process Regression

Gaussian Process regression was done using a squared exponential kernel (`RBF()`) the GPy^2^ Python package called in `Julia` through `PyCall.jl`^3^. GP hyperparameters were optimised using the `optimize_restarts` attribute of the model with 3, 5 or 15 restarts. In certain cases (stochastic *in silico* gene expression data), the length-scale parameter of the RBF kernel was fixed prior to optimisation.

## D.3 Parameter Optimisation

ODE parameter optimisation was done using the Nelder Mead [43] (`NelderMead()`) algorithm from `Optim.jl`^4^. In case of constrained parameter optimisation, this was carried out using the `Fminbox{NelderMead}()` method from the same package.

GP hyperparameter optimisation for the non-parametric inference approach was done using `GPy` as described in the previous section.

## Footnotes

1 https://github.com/JuliaDiffEq

2 https://github.com/SheffieldML/GPy

3 https://github.com/JuliaPy/PyCall.jl

4 https://github.com/JuliaNLSolvers/Optim.jl

